# Multimodal imaging reveals surface degradation of voice prosthesis implants resulting from colonisation by polymicrobial biofilms

**DOI:** 10.64898/2026.09.07.749877

**Authors:** Louise MacGowan, Liam Rooney, Pamela Thompson, Sarah De Blieck, Jan Stanier, Catriona Douglas, Gail McConnell

## Abstract

Total laryngectomy is performed for advanced laryngeal and hypopharyngeal cancers. Following the surgery, the insertion of indwelling silicone-based voice prosthesis (VP) implants is considered the gold standard for speech rehabilitation. However, early VP failure due to biofilm formation and invasion of the VP surface can lead to routine use of nystatin and the need for frequent valve changes. Despite these burdens, the composition and impact of colonising pathogens on VP integrity are poorly understood. We have developed a rapid pipeline which permitted the transfer, fixation, staining, and imaging of explanted VPs from patients within 90 minutes of sample collection. Using this pipeline, we demonstrate a multimodal reflection contrast and fluorescence confocal laser scanning microscopy technique to simultaneously map the VP surface and visualise the biofilms *in situ*, providing new insights into the role of pathogens on VP failure. Our results reveal that complex fungal and bacterial biofilms form polymicrobial biofilms on the VP surface. We demonstrate that biofilms not only reside on the VP surface but can invade the device, releasing silicone particles that are visible within the volume of the biofilms and are separate from the VP surface. Our findings highlight the complexity of the biofilms that form on speech valves; complex polymicrobial biofilms require more advanced treatment than what is conventionally used. Work should focus both on the patient factors that may influence this and the material properties of the valve.

## Introduction

Surgical removal of the larynx (total laryngectomy) is performed for advanced laryngeal and hypopharyngeal cancer [1]. Globally, over 200,000 new cases of laryngeal and hypopharyngeal cancer are diagnosed each year [2]. Following total laryngectomy, patients can experience loss of voice, loss of sense of smell, and difficulties swallowing. The gold-standard for restoring voice in a laryngectomy patient is trachea-oesophageal puncture and insertion of an in-dwelling silicone-based voice prosthesis (VP) [3, 4]. This unidirectional valve implant, the geometry of which is shown in Figure 1A, allows air to flow from the lungs into the oesophagus whilst preventing aspiration of food and liquid into the trachea [5]. Maintaining the integrity of the valve, shaft and anchoring flanges is integral to the function of the VP; failure of the device, exposes the patient to risks including aspiration pneumonia [6, 7].

**Figure 1:**
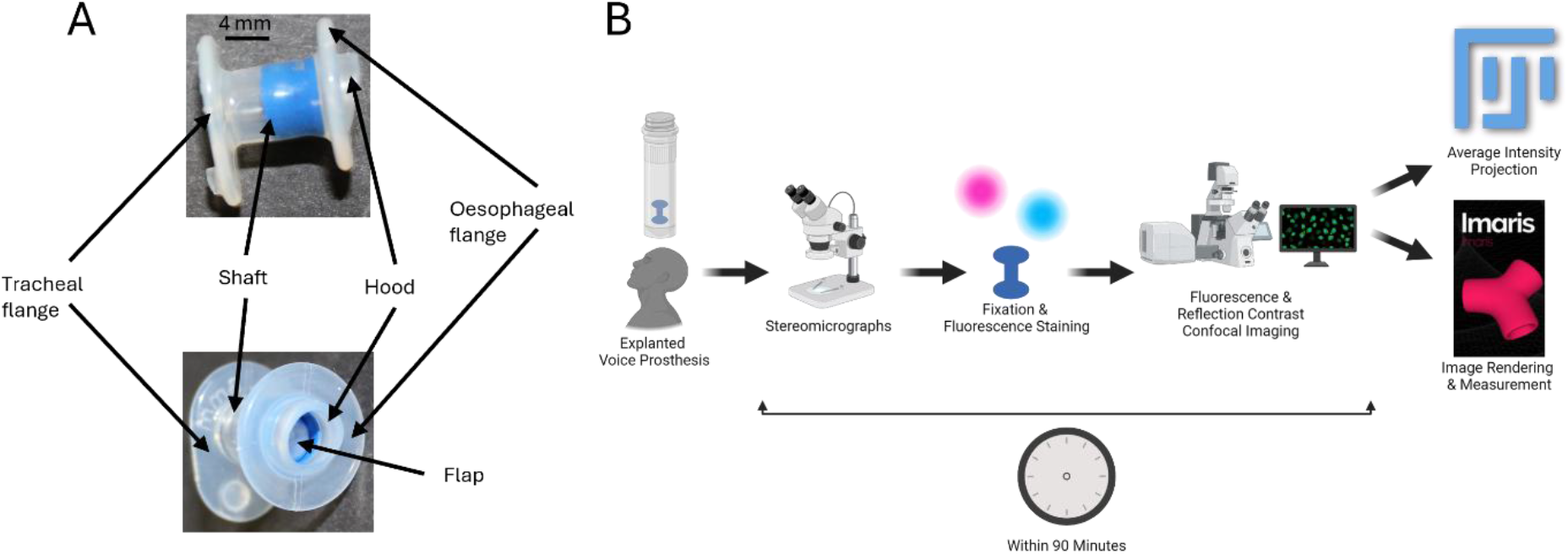
Overview of a VP and experimental pipeline. **(A)** Images indicating the location of the oesophageal flange, tracheal flange, valve hood, flap, and shaft. **(B)** The experimental pipeline from explantation of VPs at the outpatient clinic through to data analysis.

Despite their transformative impact on communication and quality of life, these devices have limited durability and can require replacement within two to three weeks [8–10]. A primary factor causing failure is the colonisation of prosthetic surfaces by biofilms, which are structured microbial communities encased in an extracellular polymeric matrix that promote surface adhesion [11–14]. Biofilms confer resilience against environmental stressors, host immune responses, and antimicrobial agents, making them recalcitrant to many treatments [15–17].

In the context of VPs, biofilm formation is predominantly attributed to fungal species like *Candida albicans* and bacterial species such as *Staphylococcus aureus* and *Lactobacillus gasseri* [18–22]. The warm, moist environment of the pharyngoesophageal region provides ideal conditions for these microorganisms to thrive. The biofilm matrix facilitates adhesion to the prosthetic surface and protects the embedded microbes from mechanical cleaning and antimicrobial interventions. These biofilms often form complex polymicrobial ecosystems, with fungal and bacterial interactions enhancing colonisation and resistance to treatment [23].

The consequences of biofilm formation on VPs can be significant: colonising pathogens can obstruct the valve mechanism, resulting in respiration of fluids through the valve and difficulty producing tracheoesophageal voice [24]. Additionally, beyond mechanical failures, biofilms pose a risk of systemic and localised infections, potentially exacerbating patient morbidity [25, 26].

Efforts to combat biofilm-associated failures have highlighted the need for preventative strategies rather than reactive measures. Surface modifications, such as altering hydrophobicity or adding antifouling coatings, have shown promise in reducing initial microbial adhesion [7, 20, 27]. However, these surface functionalised VPs are scarcely used in the clinic due to their high cost. Additionally, the inherent resilience of biofilms means that once established, they are very difficult to eradicate.

Analysis of biofilms on VPs has been carried out using different imaging methods. Scanning electron microscopy has previously been used to visualise VPs that have been reported as failing through leakage by patients, however the area that could be studied is very small and results were not statistically significant [28]. Fluorescence *in situ* hybridisation (FISH) has been used together with confocal laser scanning microscopy (CLSM) to study a larger volume of the VP and the colonising biofilm, but no information on the integrity of the VP was possible using this approach [20].

To help better understand the role of pathogens in VP failure, we have developed a multimodal reflection contrast and fluorescence confocal laser scanning microscopy technique for simultaneous imaging of the biofilms colonising VPs and mapping of the VP surface and degradation resulting from pathogen invasion. Our non-destructive method can be rapidly applied to explanted VPs, with our pipeline providing both qualitative and quantitative measures of biomass and measurements that can be used to assess the degree of degradation of VPs. This method offers the possibility of providing targeted therapeutics for patients experiencing VP failure, thus improving the quality of life of their patient cohort.

## Materials and Methods

### Experimental pipeline

A pipeline was established to facilitate the project, which was conducted as a single-blind study, to minimise bias in experimental procedure and data analysis. Over a six-week period 21 VPs were explanted due to device failure and transported from the outpatient clinic at the hospital (Glasgow, UK) to the University of Strathclyde (Glasgow, UK). The experimental pipeline for fixation, staining and imaging of biofilms in VPs was optimised using n=5 VPs that were excluded in our image analyses. 16 valves were used for the experimental research after optimisation of techniques.

A naïve Provox Vega VP (Atos Medical, Sweden) is shown in Figure 1A, highlighting the location of the integral parts of the device. The final experimental pipeline (Figure 1B) permits for data acquisition and analysis within 90 minutes of collection of the VP specimens.

### Stereomicrographs of explanted VPs

Upon receipt, stereomicrographs of the VP surface were acquired to inspect them prior to fixation, staining and imaging by multimodal CLSM, and to hallmark surface features and assess biofilm burden. Stereomicrographs of the tracheal and oesophageal flanges of each VP was acquired at 0.75× magnification, providing a 12 mm by 12 mm field of view. The images were acquired using a SMZ1500 Stereoscopic Zoom Microscope (Nikon, Japan) coupled to a 12-bit CMOS camera (DFK-33UX250; The Imaging Source, USA) controlled by IC Capture image acquisition software (version 2.4; The Imaging Source, USA).

### Fixation and fluorescent staining of biofilms

Gram-positive bacteria and fungi were selectively stained for *in situ* visualisation of biofilms. Prostheses were fixed by submerging in 3 ml of 4% (w/v) paraformaldehyde, washed three times with 1 ml of phosphate buffered saline (PBS) solution, quenched by submerging in 3 ml of 50 mM ammonium chloride, and then washed twice with 1 ml of PBS. The staining process was performed sequentially: we selectively stained Gram-positive bacteria using 1 ml of 140 nM vancomycin-BODIPY FL (ThermoFisher Scientific, USA), incubated at 37 °C for 15 minutes, and washed twice in 1 ml of PBS to remove any unbound dye. Then, fungi were fluorescently stained using 1 ml of 10 µM calcofluor white (Honeywell, USA), incubated at 37 °C for 15 minutes, and washed twice in 1 ml of PBS.

### Confocal laser scanning microscopy of VPs

Voice prosthetics were mounted on a confocal laser scanning microscope (CLSM) to visualise the biofilms *in situ* and map the VP surface. Explanted VPs were mounted in 1 ml of PBS in a 35 mm diameter polymer coverslip high walled µ-dish (ibidi, Germany). The naïve VP (Figure 1) was mounted in the same orientations as described above for imaging and the surface of the naïve VP was inspected. An Olympus IX81 inverted microscope coupled to a FluoView FV1000 confocal laser scanning unit (Olympus, Japan) was used to image the VPs. The microscope was equipped with an UPLSAPO 10×/0.4 numerical aperture (NA) objective lens (Olympus, Japan). Excitation wavelengths of 405 nm and 488 nm were used to excite calcofluor white and vancomycin-BODIPY FL, respectively. A 633 nm laser was used for reflection contrast imaging of the VP surface. The 405 nm laser wavelength line was provided by a diode laser (model no. GLG3135; Showa Optics, Japan), the 488 nm laser line was provided by an argon laser (model no. GLG3135, Showa Optics Japan), and the 633 nm laser line was provided by helium-neon laser (model no. GLG3135, Showa Optics, Japan).

The microscope was configured for epifluorescence detection using a 405/488/633 nm dichroic mirror and the detection bandwidths for fluorescence from calcofluor white, vancomycin-BODIPY FL and reflection from the VP surface were set to 420-470 nm, 535-600 nm and 630-640nm, respectively. The resolution of the microscope is given by wavelength (λ) and NA. The theoretically calculated lateral resolution, given by r_lat_ = 0.61λ/NA, at λ=530 nm is 808 nm, and the axial resolution is 2λ/NA^2^ is 6.6 µm. All images were acquired with at least two pixels per resolution unit of the microscope to satisfy the Nyquist-Shannon sampling criterion. Confocal z-stacks of the oesophageal and tracheal flanges of 16 VPs were acquired, using a z-step size of 1.2 µm. Data were saved in the proprietary. OIB format and were exported to OME TIFF for analysis.

### Image processing

Confocal z-series were processed using FIJI (v1.54k) [25] and image data were presented as average intensity z-projections of each image stack. The images were linearly contrast adjusted for presentation purposes.

The mounted specimens were tilted relative to the optical axis because of the non-planar geometry of the VP, and so the position of the specimen led to reflection from the coverslip in the acquired images. For presentation purposes, coverslip reflection in the images was minimised by using the *‘Crop 3D’* and *‘Add Slices’* functions in Imaris (v9.8.0; Oxford Instruments, UK). The reflection channel data was cropped using a 1.2 µm step size in the z-direction. This reduced the z-dimension of the reflection data but removed the coverslip reflections which confounded visualisation of the data. The cropped slices were replaced with empty (i.e. dark) slices to equalise the volume of the reflection channel to the fluorescence channels for further processing. Overlays of the reflection and fluorescence channels were generated in Imaris.

### Image analysis of biofilms and silicone particles

Image analysis was implemented to visualise and quantify the fungi and Gram-positive bacteria in the biofilms. To quantify the volumes of fungi and Gram-positive bacteria surface analysis was performed in Imaris. The thresholding parameters were adjusted to exclude background signal from the biofilm volume and the contrast was linearly adjusted for visualisation purposes. The thresholding parameter for *‘split touching objects’* was limited to 5 µm because of the axial resolution of the microscope. Selecting a smaller object size resulted in single objects being counted twice. To calculate the relative volumes of fungi and Gram-positive bacteria within the total biofilm volume observed in the z-stacks, normalisation was performed in Excel. The data were normalised by presenting either the total fungal or total bacterial volume as a fraction of the total biofilm volume (i.e., fungi plus bacteria). The relative volumes of fungi and bacteria were plotted using Prism (v8.0.2, GraphPad, USA).

Surface analysis was performed for the reflection contrast channel to quantify the volume of silicone particles in the biofilms on VPs. The analysis was revealed there were silicone particles visible within the biofilm mass on seven VPs. For each VP, three regions of interest (ROIs) distal to the VP surface measuring 100 µm x 100 µm x 100 µm were selected to exclude the VP surface and reflection from the coverslip. The same procedure described above was applied to quantify the volume of sloughed silicone in the reflection channel. Particle sizes were plotted using SuperPlotsOfData [29].

## Results

### Non-uniform surface topography of naïve voice prostheses

Stereomicrographs of the oesophageal and tracheal flanges of the naïve VP are shown in Figures 2A and 2B respectively. A yellow arrow indicates the valve flap, which opens during speech and otherwise remains closed to prevent leakage of food and saliva into the trachea [11]. The blue fluoroplastic tube inside the valve shaft prevents the valve flap from opening in the tracheal direction. Numerals formed on the VP surface denote the length and diameter of the valve shaft. A magnified region, designated by the black square in Figure 2A, was imaged using a CLSM configured in reflection mode to assess the surface profile of the VP, and a representative image of a VP is shown as an average intensity z-projection in Figure 2C. Imperfections on the VP surface such as scratches and pits can be observed in 2C, with indentations giving rise to an uneven surface topography at the microscopic scale. Orthogonal views of the 3D CLSM data are presented for x, z (Figure 2D) and y, z viewpoints (Figure 2E). Figures 2D and 2E show the tilt of the specimen relative to the optical axis of the microscope, which results from the curved geometry of the hood around the valve opening causing the VP to sit at an angle to the coverslip.

**Figure 2:**
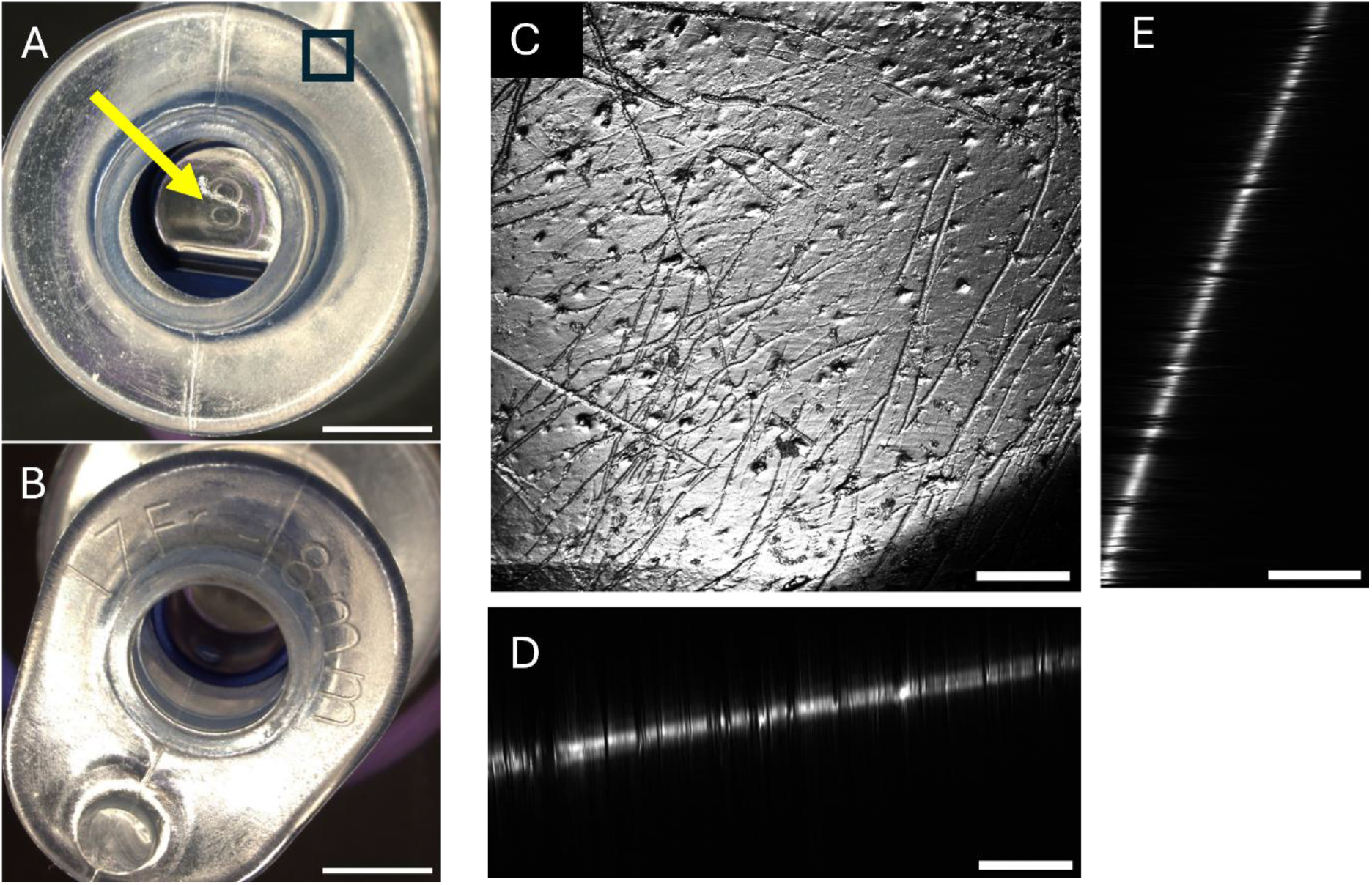
Surface topography of a naïve VP. Stereomicrographs presenting the oesophageal **(A)** and tracheal flange **(B)** of a naïve VP, where the yellow arrow indicates the valve flap. **(C)** An average intensity z-projection of a 3D CLSM acquisition of the region noted by the black square presented in (A). **(D)** and **(E)** show the orthogonal views of the z-projection shown in (C). Due to the geometry of the VP the surface does not sit perpendicular to the optical axis. Scale bars for A-B = 300 µm, scale bars for C-E = 200 µm.

### Biofilm surface colonisation of explanted VPs

The oesophageal flange and tracheal flange of 16 explanted VPs imaged using a stereomicroscope are shown in Figure 3. The devices shown are manufactured by Atos Medical (Sweden) and InHealth Technologies (USA). Visual inspection of the stereomicrographs indicated that the biofilm burden was predominantly located on the oesophageal flange and valve flap of the VPs. However, the degree, location, colour, and form of colonisation were observed to differ across the patient cohort. The blue residue visible on VPs 14, 15 and 16 was a result of residual blue food colouring used in the clinic to test for valve leakage before explantation. The indwelling time of the VPs shown was between 12 and 347 days, with a median indwelling time of 88 days and IQR 136.5 days. There was no direct connection between the observed coverage of the biofilm versus the indwelling time. For example, the indwelling time of VP 12 was 86 days, and the indwelling time of VP 15 was 81 days. Biofilms which colonised the oesophageal flange and valve flap of VP 15 cover most of the surface, whereas on the oesophageal side of VP 12 the biofilms can be observed around the opening to the valve shaft and covering only a small portion of the oesophageal flange. Biofilms appeared to cover much of the oesophageal flange of six failed VPs, whereas the bioburden on ten failed VPs covered a much smaller proportion of the oesophageal flange. As such, we observed no direct connection between extent of device colonisation and VP failure.

**Figure 3:**
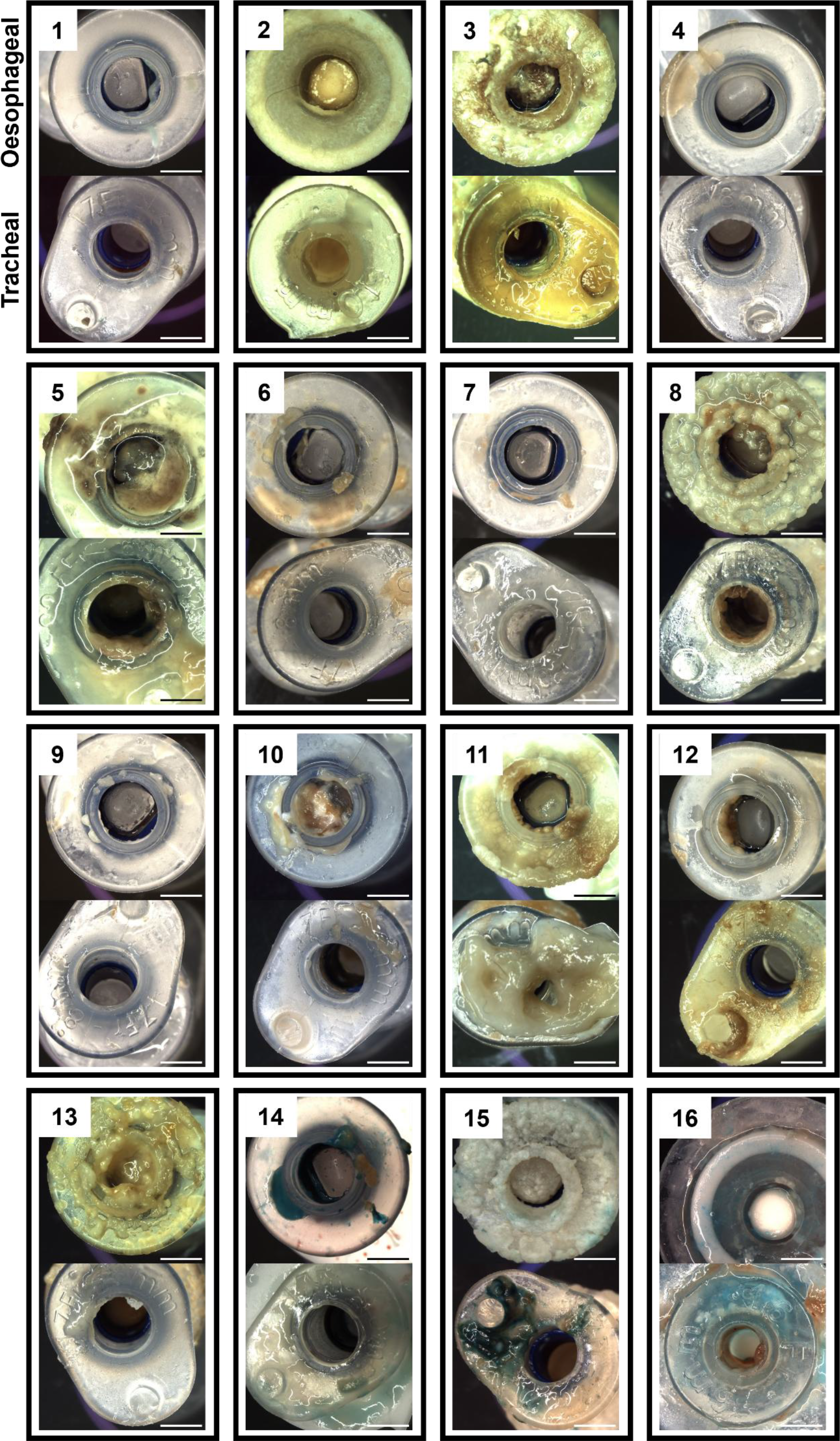
Stereomicrographs of explanted VPs illustrating biofilm colonisation. An overview of the biofilm colonisation of each VP in the patient cohort taken forward for multimodal imaging. Top panels show stereomicrographs of the oesophageal flange and the bottom panels show the tracheal flange. Scale bars = 300 µm.

### Architecture of biofilms on explanted VPs

Fluorescence and reflection contrast CLSM images of explanted VPs reveal varying degrees of colonisation of the surface by complex polymicrobial biofilms, but all imaged ROIs showed degradation of the silicone surface relative to the naïve device. Figure 4 presents 3D images of the oesophageal flange of three representative VPs with different bacterial burdens. Average intensity z-projections for all VPs analysed are provided in Supplementary Figure 1. Figure 4 panels (A-C), (D-F), and (G-I) correspond to VPs 12, 10 and 3 shown in Figure 3, respectively. The top row shows a composite merge of fluorescently stained fungi, Gram-positive bacteria, and surface reflection. The central row shows fluorescence from fungi stained with calcofluor white, and the bottom row shows fluorescence from the Gram-positive bacteria stained with vancomycin BODIPY-FL. Figure 4(A-C) show discrete biofilms of Gram-positive bacteria with two fungal microcolonies, indicated by yellow arrows. The composition of biofilms present on different VPs was heterogenous and most biofilms observed were polymicrobial. Biofilms comprising fungal mycelia with Gram-positive bacteria distributed throughout the biofilm were observed, as shown in Figure 4(D-F). Biofilms comprising fungi and discrete microcolonies of Gram-positive bacteria within the biofilm were also observed, as shown in Figure 4(G-I). Average intensity projections and 3D reconstructions of these biofilms are shown in Supplementary Figure 2. Objectively, biofilm morphology was not consistent between VPs and did not correspond to bioburden. Scores observed on the surface of the naive VP (Figure 2C) were not observed on explanted VPs indicating erosion of the silicone whilst *in situ*, as shown in Figures 4A and 4D.

**Figure 4:**
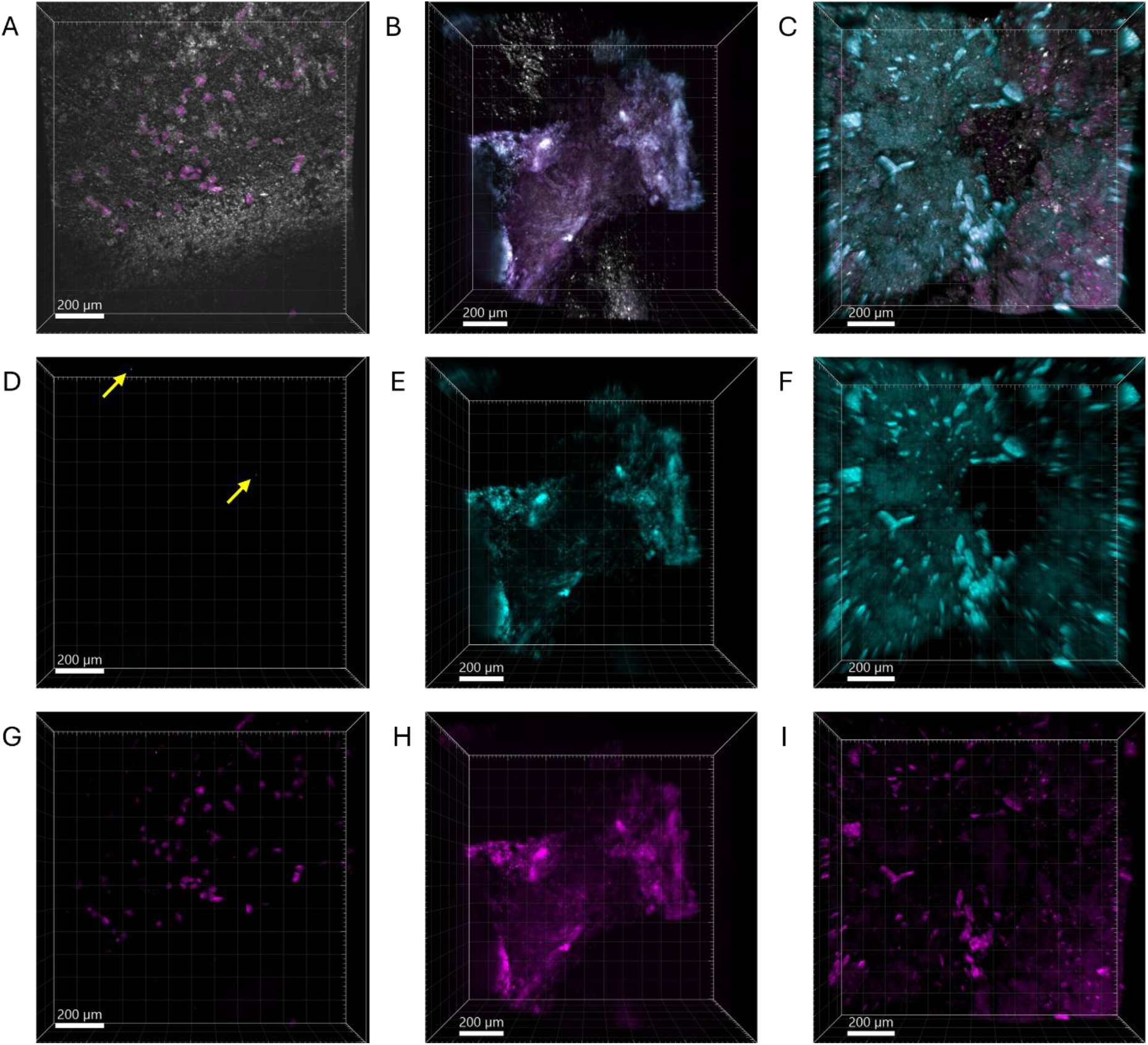
Surface colonisation patterns of VPs. Fluorescence and reflection contrast 3D CLSM images of biofilms on the oesophageal flange of explanted VPs. From left to right, the images show VPs with increasing bioburden. Fungi are presented in cyan, Gram-positive bacteria in magenta, and surface reflection in grey. Scale bars = 200 µm. **A, D, E**: Overlays of fungi, Gram-positive bacteria and surface reflection. **B, E, H**: Fungi. **D, F, I**: Gram-positive bacteria. The yellow arrows in (B) indicate microcolonies of fungi and the white boxes indicate a magnified ROI.

### Biofilm architecture and surface degradation

Polymicrobial biofilms up to 300 µm thick were observed on explanted VPs (Figure 5A). The biofilm contours corresponded to the topography of the VP surface. Biofilms growing on the outer edge of the oesophageal flange (Figure 5B), or over the valve hood (Figure 5D), exhibited reflection from silicone distal to the VP surface. Figure 5E illustrates a biofilm with regions of low fluorescence and hyphae extending throughout the volume of the biofilm. No silicone particles were present in the biofilm volume (Figure 5F). This implies that the mechanism of adhesion to and invasion of the silicone differs from that of the biofilms in 5B and 5D. In Figure 5C, the yellow arrow indicates a volume of silicone distal to the VP surface. The reflection did not coincide with fluorescence from the biofilm, indicating that silicone had been released from the VP surface. Silicone particles, distal to the VP surface, exhibited a distribution with the same morphology as the biofilm contours (Figure 5B and 5D), thus demonstrating that biofilms degrade VPs by drawing silicone particles from the VP surface. 3D animations for Figures 5A-F are provided in Supplementary Movies 1, 2, and 3.

**Figure 5:**
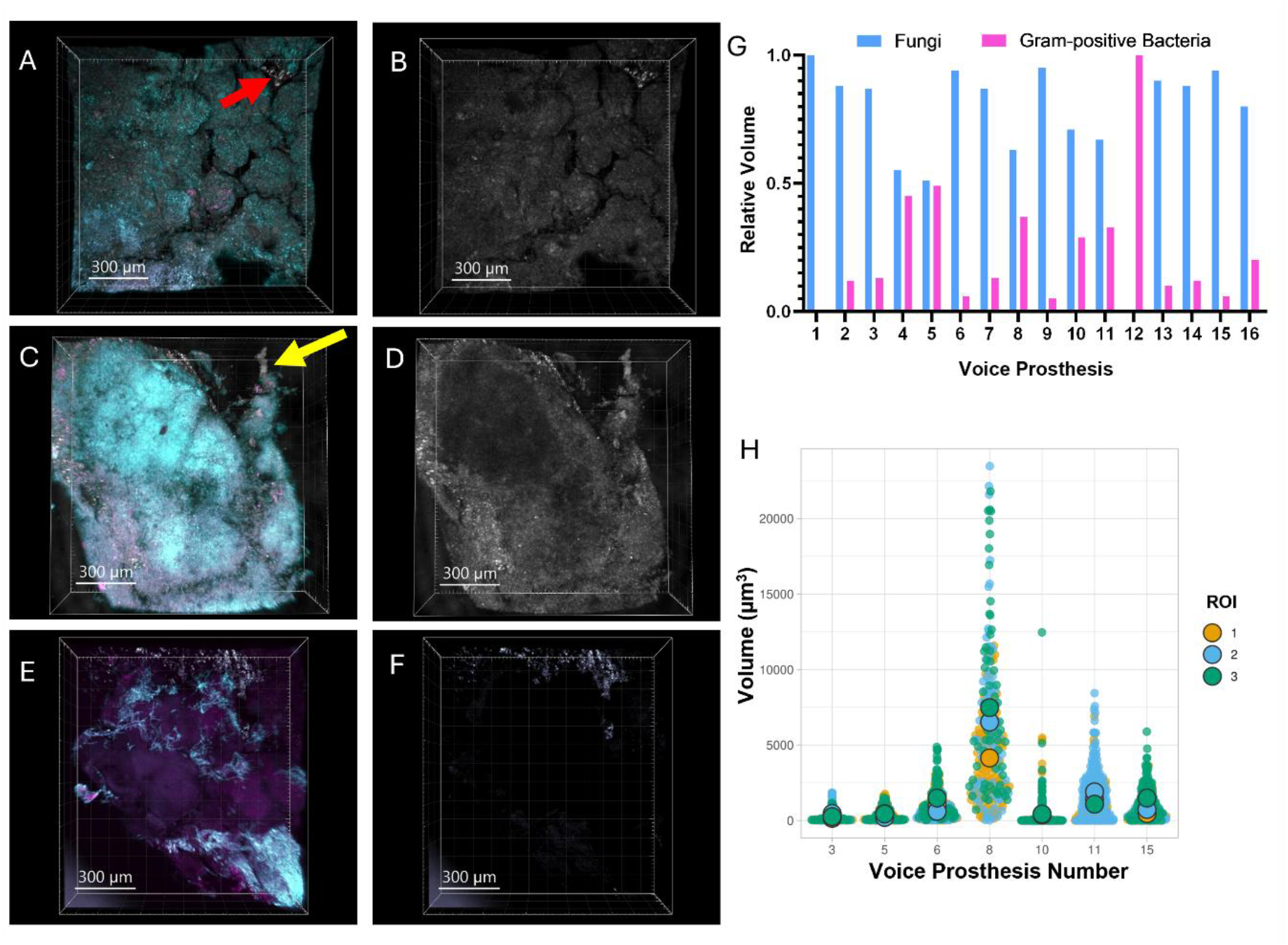
Silicone particles in VPs resulting from colonizing pathogens. 3D CLSM images of fluorescently stained fungi (cyan), Gram-positive bacteria (magenta), and surface reflection (grey). The images are of VPs 15 (A, B), 8 (C, D), and 13 (E, F). **(A, D, and E)** show overlays of fungi, Gram-positive bacteria, and surface reflection. The red and yellow arrows in (A and C) indicate sloughed regions of the VP surface. **(B, D, and F)** Surface reflection. **(G)** Relative volumes of fungi and Gram-positive bacteria in the biofilm volume. Animations of the 3D images are provided in Supplementary Movies 1, 2, and 3. **(H)** The size of silicone particles in three replicate ROIs for each VP are shown in green, blue, and yellow. The size of the black outlined circles represents the number (N) of particles in each ROI and the position indicates the mean particle volume. Each ROI was a cube of volume 100 µm x 100 µm x 100 µm.

Normalised data from the surface analysis of fungi and Gram-positive fluorescence volume (Figure 5G) shows that fungi generally occupy a larger volume of the biofilm than Gram-positive bacteria. For VPs 4 and 5 fungi and Gram-positives occupied similar volumes of the biofilms. On VP 4 biofilms cover less than 25% of the total area of the oesophageal flange and there is no biofilm visible on the valve flap. On VP 5, biofilms cover half of the valve flap and have colonised more than one quarter of the oesophageal flange (Figure 2). The indwelling times for these VPs was 324 days and 201 days, respectively. The in-dwelling time for VPs 1 and 12 was 49 days and 86 days, respectively, and the bioburden on these VPs was low. On VP 1 the biofilm was entirely fungal but on VP 12 the biofilms were Gram-positive with spatially disparate fungal microcolonies. The biofilm volume was predominantly fungal on most of the VPs; however, the volume analysis revealed there is a significant number of viable Gram-positive bacterial cells present in the biofilms.

Results for the surface analysis of silicone particles on seven VPs, which exhibited silicone particles within the biofilms, are shown in Figure 5H. VPs which did not exhibit active degradation by biofilms were excluded from the silicone particle analysis. Three replicate ROIs for each VP are plotted in green, blue, and yellow. The number of particles contained in the ROIs ranged between two and 441. The mean volume of particles in all ROIs, except for VP 8, was less than 2000 µm^3^, with a mean diameter of approximately 13 µm. For VP 8 there was a large spread of particles with volumes greater than the mean in all three ROIs. For VPs 3 and 5 a small spread of volumes above the mean was observed in the ROIs. The range of particle volumes distributed throughout the biofilms was inhomogeneous.

## Discussion

Our silicone particle analysis demonstrated that biofilms actively damaged the VP surface of seven devices included in this study. Silicone particles were not observed in all biofilms (Figure 5 E and F), suggesting that surface degradation may be dependent on the composition of the biofilm. Degradation of the VP surface has been demonstrated previously, following removal of the biofilms or dehydration of biofilms after sample preparation for scanning electron microscopy [21, 37]. CLSM demonstrates that diverse polymicrobial biofilms containing a significant volume of Gram-positive bacteria colonise VPs. In some instances, Gram-positive bacteria and fungi were evenly distributed throughout the biofilm and, in others, Gram-positive bacteria formed discrete microcolonies within the polymicrobial biofilm. The variation in biofilm architecture suggests that biofilm formation on VPs is not a uniform process. The location and architecture of biofilms may contribute to VP failure. Colonies located around the valve flap may prevent valve opening or closure and the extent of Gram-positive bacteria may impact on the efficacy of antifungals.

The naïve VPs had an uneven surface with microscopic features such as scratches and pits, which has been reported using scanning electron microscopy and atomic force microscopy [30]. These microstructures were observed on both the tracheal and oesophageal flanges of the VPs. Numerals moulded onto the tracheal flange and valve flap during the manufacturing process further contributed to the roughness of the surface. Newly implanted VPs may be predisposed to colonisation because biofilms form more readily on rough and damaged surfaces [31, 32]. These surface irregularities may provide accessible sites on which bacteria and fungi can adhere.

The surface of explanted prosthesis (Figure 4A) relative to the naïve VP (Figure 2C) was observed to be altered on all VPs. Scratches observed on the naïve VP were not present on the surface of explanted VPs suggesting erosion of the surface of explanted devices. This may be due to the passage of food over the VP, enzymes produced in the surrounding tissue, or colonisation and subsequent shedding of biofilms from the VP surface. Silicone particles released from the VP could possibly provide vehicles for biofilms to migrate into the patient’s aerodigestive tract. The effects of this on the patient is unknown. They may also provoke an inflammatory response in surrounding tissues, potentially leading to irritation, discomfort, or tissue damage at the site of the prosthesis [38, 39].

Biofilm burden was predominantly observed on the oesophageal flange and valve flap, with variability in the extent of biofilm burden between VPs. The observed macroscopic extent of biofilm burden was not indicative of device failure. Differences in the visual appearance, colour and shape, of biofilm reveals that biofilm morphology varied between VPs (Figure 3). This indicates that other factors, such as the unique yet transient microbiome of the patient may play a significant role in the failure rate of VPs [33–35]. Additional factors, such as lifestyle and diet, are also likely to influence the device lifetime [36].

Any Gram-negative bacteria present on the VP surface may also contribute to the volume of the biofilm, and it would be interesting to understand their role in device failure by modifying our pipeline to incorporate additional fluorescent stains to label anoxic cell populations, for example. This was not possible on our microscope, but advanced spectral unmixing algorithms offer the possibility of imaging further channels using our multimodal approach.

The failure of speech valves is complex, and reasons may be multifactorial. Our study has thus far been localised to the surface of the VP, but it would be interesting to adapt our pipeline for the structural study of the VP flap. Silicone particles caused by invading pathogens may accumulate within the valve mechanism, obstructing airflow, increasing resistance, and contributing to valve leakage. This would compromise the patient’s ability to produce sound, and may also impact on swallowing, necessitating more frequent device replacements. For this work, a microscope objective lens with a modest magnification, high NA, and long working distance would be required, representing an opportunity for development in new optical technology.

### Conclusion

Fluorescence and reflection contrast CLSM can be used to assess the fungal and Gram-positive bacterial composition of biofilms on VPs, without the need for scraping or dissection of the device. Significant volumes of Gram-positive bacteria are present in these complex polymicrobial biofilms and are likely to be implicated in VP failure. Prophylactic use of antifungals to combat *Candida* biofilms may not be the most appropriate treatment for all patients. Furthermore, degradation of the VP may lead to silicone particles entering the patients aerodigestive tract, which could potentially have implications for the patient.

Better understanding of biofilm formation on VPs could allow for more targeted drug delivery and improved device design. Development of new materials or surface coatings that reduce surface roughness and inhibit biofilm formation is required to improve VP design. Thus, reducing the use of systemic antimicrobials or topical application of prophylactic antimicrobials. This study suggests that in future there may be an additional step to consider i.e. whether there is also a more personalised and bespoke-method of managing individual patterns of degradations.

## Ethics

Ethical approval for this study was obtained from the NHS Research Scotland (NRS) Biorepository Network, reference number TR 1034. The research utilised anonymised clinical data under the NRS Biorepository Network guidelines. All samples were collected with informed consent under protocols approved by the NHS Greater Glasgow and Clyde Cadicott Gaurdian.

## Acknowledgements

Figure 1 was created in BioRender. MacGowan, L. (2026) https://BioRender.com/lu9yew1 under licence number VX28IB2565.

## CRediT authorship contribution statement

**Louise MacGowan:** Writing – original draft, Visualization, Validation, Investigation, Formal analysis. **Liam Rooney:** Writing – original draft, Conceptualization, Methodology, Visualisation, Validation, Investigation, Formal analysis. **Pamela Thompson:** Resources, Writing – review & editing. **Sarah De Blieck:** Resources, Writing – review & editing. **Jan Stanier:** Writing – review & editing. **Catriona Douglas:** Conceptualization, Supervision, Funding, Writing – review & editing. **Gail McConnel:** Conceptualization, Methodology, Validation, Supervision, Funding, Writing – original draft.

## Data Availability

The experimental data generated and analysed in this study is available from the University of Strathclyde Pure repository DOI: https://doi.org/10.15129/49f04acb-6bee-43f1-ba1a-967c4a9bc12f

## Appendix A. Supplementary data

**Supplementary Figure 1: Biofilms on explanted VPs.** Average intensity projections of ROIs of all VPs carried forward for experimental research. Fluorescently labelled Gram-positive bacteria (magenta), fungi (cyan), and reflection from the VP surface (grey).

**Supplementary Figure 2: Composition of biofilms on explanted VPs. A** - **C** are 3D reconstructions**. D** - **F** are average intensity projections of the same regions as shown in A - C. Fluorescently labelled Gram-positive bacteria (magenta), fungi (cyan), and reflection from the VP surface (grey).

**Movie 1**

**Movie 2**

**Movie 3**

