## Supplementary figures and images for "Multimodal imaging reveals surface degradation of voice prosthesis implants resulting from colonisation by polymicrobial biofilms"

### Supplementary Figure 1

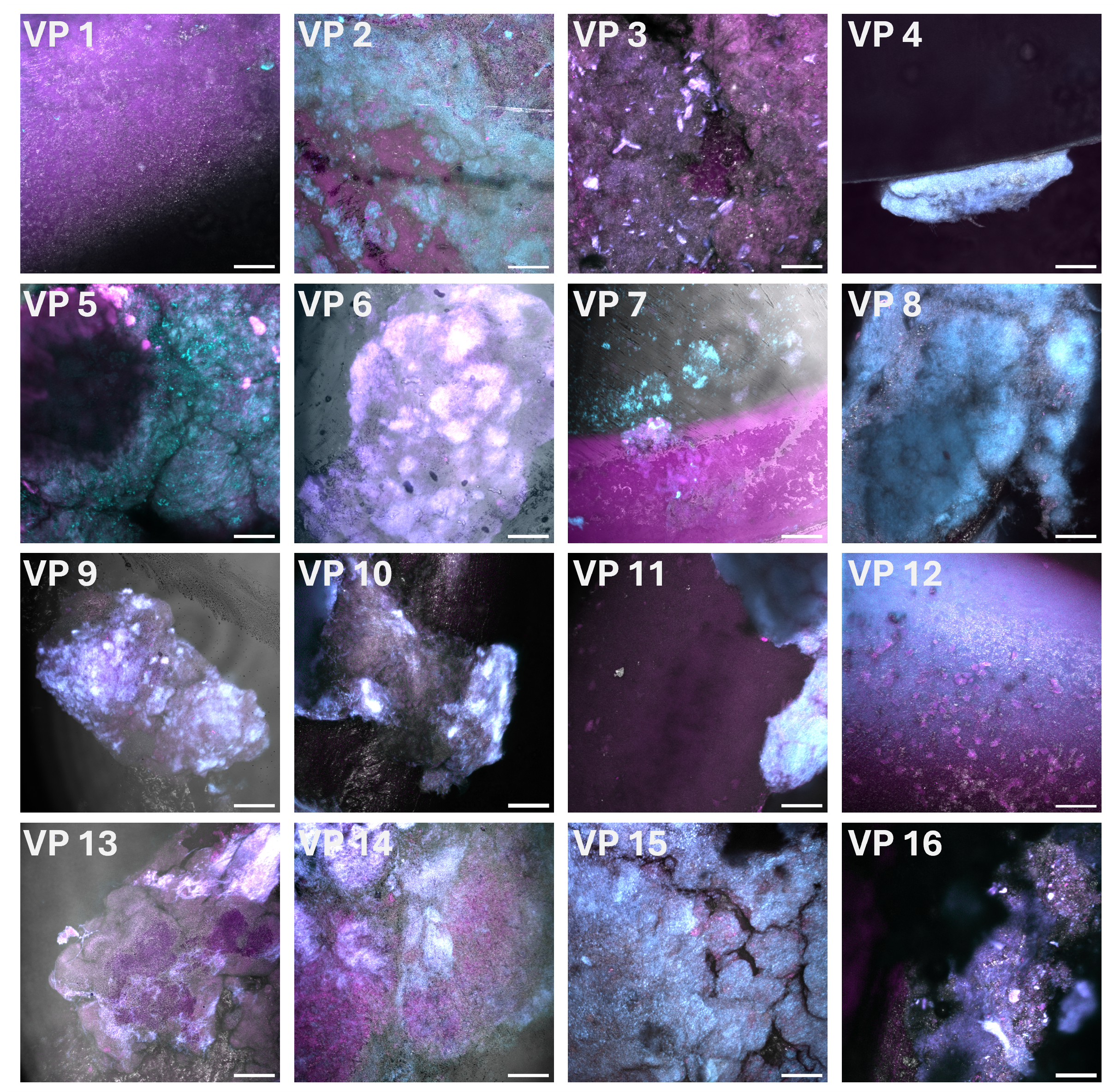

### Supplementary Figure 2

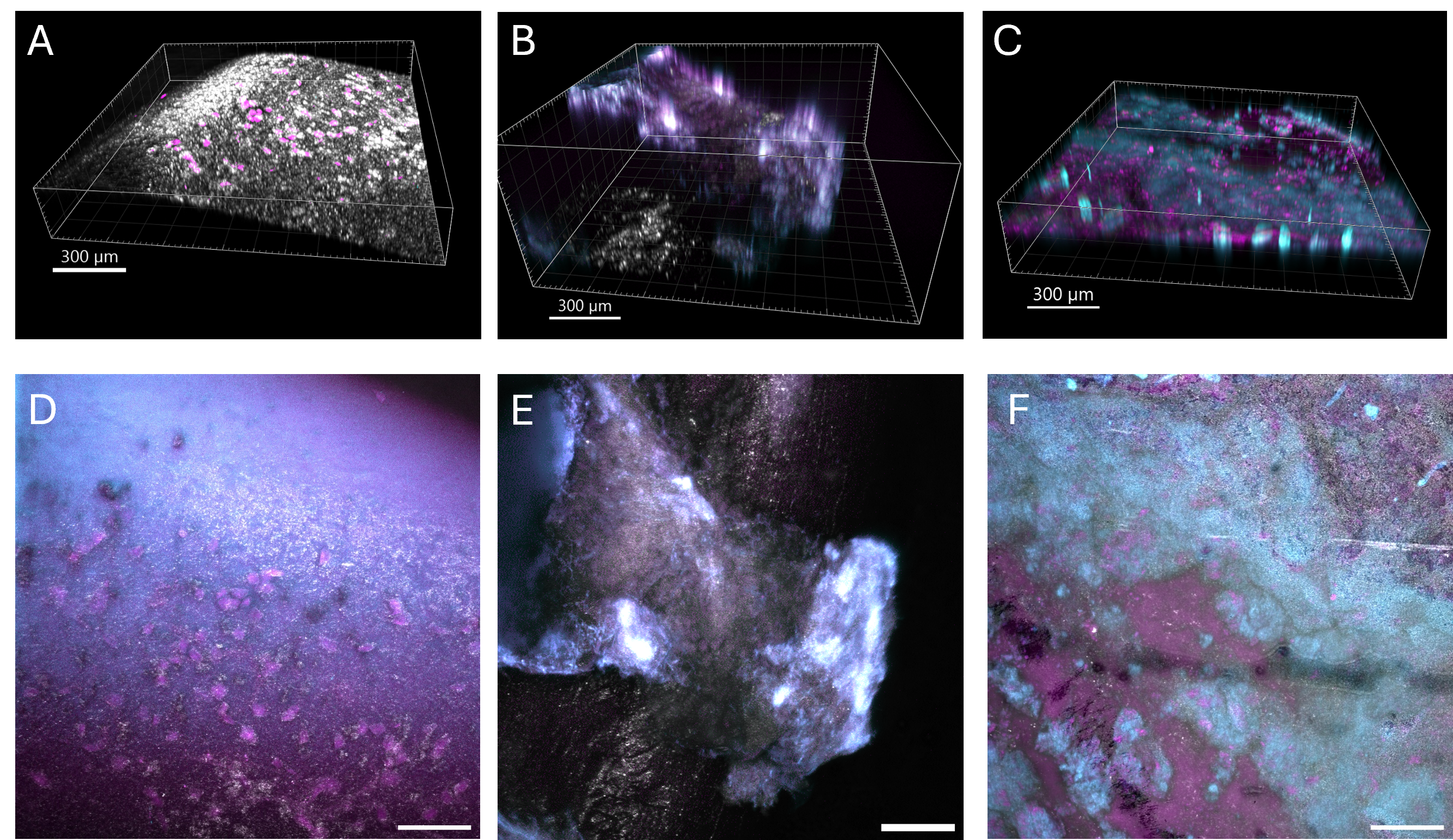
